# The evolution of a beneficial association between an animal and a microbial community

**DOI:** 10.1101/357657

**Authors:** D. Rebar, H. C. Leggett, S.M.L. Aspinall, A. Duarte, R.M. Kilner

**Affiliations:** Department of Zoology, University of Cambridge, Downing Street, Cambridge, CB2 3EJ; Department of Biological Sciences, Emporia State University, 1 Kellogg Circle, Emporia, KS 66801; College of Life and Environmental Sciences, University of Exeter, Penryn Campus, Penryn, Cornwall, TR10 9FE

**Author notes:** joint first author and corresponding author.

## Abstract

Animals are now known to be intimately associated with microbial communities, some of which enhance animal fitness. Yet relatively little is known about how these beneficial associations initially arose. We investigated this problem with an experiment on burying beetles, *Nicrophorus vespilloides*, which breed on the body of a small dead vertebrate. We found that burying beetles breeding on germ-free mice produced smaller larvae, with lower fitness, than those breeding on conventional germ-laden mice. Thus, burying beetles gain benefits from the microbial community associated with their carrion breeding resource, because they lose fitness when this community is removed experimentally. Our experiment suggests that a symbiosis between an animal and a microbial community might begin as an adaptation to the microbial ecosystem in which the animal lives, even when these microbes exist outside the animal, are transiently associated with it at each generation and are not directly transmitted from parents to offspring.

## BACKGROUND

Recent advances in sequencing technology have revealed that all animals are intimately associated with extensive microbial communities, often carried within or upon their bodies. Furthermore, whereas microbes have traditionally been regarded as antagonists that reduce animal fitness (e.g. Janzen 1977), many of the microbial communities associated with animals are now recognised to confer benefits that enhance animal fitness. Microbial communities can contribute to animal health (e.g. Flint et al 2012), for example, and perform other key services (e.g. Ezenwa & Williams 2014, Ruokolainen et al 2016). Yet relatively little is known about how these beneficial associations might have arisen in the first place.

Evolutionary theory predicts that beneficial associations between a microbial community and an animal will arise and persist when the evolutionary interests of all the parties are similarly aligned (e.g. Douglas & Werren 2016, Moran & Sloan 2015, Fisher et al 2017). These conditions are most likely to be met when the animal and the microbial community are intimately, and enduringly, associated with one another and the microbial community is transmitted from animal to animal across the generations with high fidelity. To test these predictions, a recent comparative analysis of animals and bacterial symbionts investigated which factors best explained the extent to which animal hosts are dependent on bacterial symbionts, where dependency was measured by quantifying the animal’s fitness loss when the bacteria were experimentally removed (Fisher et al 2017). This study found that animals had evolved a greater level of dependence when the bacteria performed a nutritional service for their animal hosts and when they were vertically transmitted from host parents to offspring (Fisher et al 2017). However, it is not straightforward to disentangle which of these two properties might have contributed most to the evolutionary origins of any such beneficial association. An animal-microbial community mutualism cannot evolve unless the microbes are performing a service of some sort for the animal, but nor is there the opportunity for a service to evolve unless the animal is recurrently exposed to the microbial community, which is most-readily achieved via vertical transmission.

To address this conundrum, we tested whether animals can gain fitness from a microbial community when the microbial community is not directly vertically transmitted from animal parents to offspring. Our experiments focused on the burying beetle, *Nicrophorus vespilloides*, which breeds on the body of a small dead vertebrate. Burying beetles convert the dead body into an edible carrion nest for their larvae by biting into the dead animal’s abdomen, pulling out the gut and consuming it (Duarte et al 2017). Then they bite off the fur (or feathers), roll the flesh into a ball and bury the ball below ground, all the while smearing it with fluids from the mouth and anus (Cotter & Kilner 2010). These activities increase the bacterial load on the carrion nest, partly because some of the bacteria ingested when the beetles consume the vertebrate’s skin and gut apparently pass into via the anal exudates and back onto the carrion nest (Duarte et al 2017).

We predicted that if burying beetles gain fitness benefits from associating with the microbial community on mouse carrion, they should lose fitness when they breed on ‘germ-free’ mice – i.e. mice whose microbial community has been experimentally eliminated. In conventional animals, different parts of the body harbour different microbes (e.g. Ding & Schloss 2014). To determine which of these communities have most influence on burying beetle fitness, we bisected all mouse cadavers to generate a head portion (to analyse effects of mouse skin and hair microbes, and any microbes in the upper gut down to the stomach) and a tail portion (to analyse any effects of the mouse lower gut microbes, as well as the skin and hair microbes). We measured beetle fitness on each type of carrion by counting and weighing the larvae produced. Previous work has shown that larval mass at this developmental stage predicts the likelihood of survival to adulthood and adult size (e.g. Lock et al 2004), which in turn, is a strong predictor of fecundity in both sexes (Jarrett et al 2017, Pascoal et al 2018).

## METHODS

### General Methods

Beetles were maintained at 20°C and on a 16:8 light:dark cycle, as described in detail elsewhere (Jarrett et al 2017). Each portion of carrion was weighed and placed in a standard burying beetle breeding box (17 × 12 × 6 cm) lined with 3 cm of MiracleGro compost, bought commercially. A pair of unrelated, virgin burying beetles *N. vespilloides* was placed on the carrion and the breeding box was sealed with a lid and placed in a dark cupboard to simulate underground conditions. Eight days after pairing, we measured beetle fitness by weighing and counting the dispersing larvae.

### Germ-free versus conventional mice

Germ-free mice are cultured for use in medical research, and have no associated microbiota. Germ-free mice (strain C57BL/6J, N=10) were bought dead and frozen from the Wellcome Trust Sanger Institute, Hinxton, Cambridgeshire UK. They were culled, stored frozen and defrosted under sterile conditions (mean mass = 31.47g). We bisected the germ-free mice with a transverse cut, generating separate head and tail portions of approximately equal mass (15.21 ± 2.70 g and 16.54 ± 2.91 g, respectively). We compared the performance of beetles breeding on germ-free mice with albino white mice that were bought dead from Livefoods Direct Ltd (Sheffield) and of similar initial mass to the germ-free mice (mean = 32.29 g, N=10). They were also stored frozen before being defrosted for use in this experiment. They were bisected in exactly the same way to generate a head and tail portion (15.48 ± 2.96 g and 16.81 ± 2.84 g, respectively). This is the standard mouse strain used for burying beetle reproduction in our laboratory. All the beetles used in our experiments derived from a population that had bred on this strain for at least the preceding 8 generations.

### Statistics

Statistical analyses were carried out using the package lme4 (Bates et al. 2015) in R v. 3.4.1 (R Core Team 2017). The initial models analysed the effects of the manipulation on brood success, brood mass, and brood size separately. Models contained the following terms: carcass treatment (germ-free versus conventional); carcass part (head versus tail); carcass treatment x carcass part; carcass mass. To analyze the effects of our manipulation on average larval mass, we first controlled for larval density since this is known to significantly influence average larval mass at dispersal (e.g. Schrader et al 2015). We achieved this statistically by pooling the data and fitting a linear regression relating the average larval mass to larval density (Table 1). We then used the residuals from that regression to compare average larval mass between carcass treatment and carcass part. This model had the following terms: carcass treatment (germ-free versus control); carcass part (head versus tail); carcass treatment x carcass part.

**Table 1.** Best-fit regression model of mean larval mass on larval density for all broods produced.

| Source | *Estimate* | *Std error* | *t* | *P* | *R*<sup>2</sup> |
| --- | --- | --- | --- | --- | --- |
| Intercept | 0.2192 | 0.0075 | **29.26** | **<0.001** | 0.67 |
| Larval density | -0.0391 | 0.0045 | **-8.66** | **<0.001** |  |

## RESULTS

We found no difference in the overall breeding success of beetles on conventional LiveFoods versus germ-free mice: 19 out of 20 pairs on germ-free mice bred successfully compared with 20 out of 20 pairs on conventional mice (GLM with binomial distribution, logit link function; *X^2^* = 0.00, *p* = 1.00). We also found that beetles in the two treatments produced a similar number of offspring (LM; F_1, 34_ = 0.14, *p* = 0.71), and that this was irrespective of the portion of the carcass on which they developed (LM; F_1, 34_ = 0.03, *p* = 0.86). Similarly, we found that the mass of the broods did not differ with respect to the type of carcass or the body part they bred upon (LM; F_1, 34_ = 1.94, *p* = 0.17 and F_1, 34_ = 0.57, *p* = 0.45, respectively).

However, we did find a marked difference in the average mass of larvae produced in the different treatments, once we had controlled for larval density on the carcass. Contrary to our predictions, individual larvae were significantly smaller when developing on germ-free mice (Fig. 1a, b, Table 2) and larvae that were raised on the tail portion of the germ-free mice were even smaller than those raised on the head portion (Fig. 1b, Table 2). In a subsequent experiment, we established that these differences are not simply due to lab-adaptation of burying beetles to the LiveFoods conventional mouse strain, independent of any effects of the microbial community (see Supplementary Material).

**Fig. 1.**
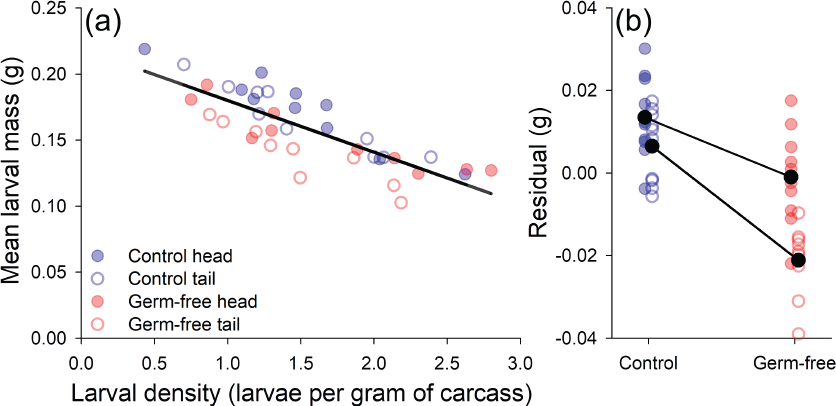
The mean larval mass of members of the brood at dispersal. A) A linear regression (*R^2^* = 0.67) describes the relationship between mean larval mass and larval density. B) The relative size of larvae at dispersal differs according to carcass type, carcass part, and their interaction.

**Table 2.** A linear model testing for the effect of carcass treatment, carcass part, and their interaction on the relative size of larvae at dispersal.

| Source | *Estimate* | *Std error* | *t* | *P* |
| --- | --- | --- | --- | --- |
| Intercept | -0.0005 | 0.0016 | -0.33 | 0.741 |
| Treatment (control) | 0.0105 | 0.0016 | **6.64** | **<0.001** |
| Part (head) | 0.0068 | 0.0016 | **4.29** | **<0.001** |
| Treatment (control) × Part (head) | -0.0033 | 0.0016 | -2.08 | **0.045** |

## DISCUSSION

We found that burying beetles gain fitness from the microbial community borne on and within the burying beetles’ carrion breeding resource. When we eliminated this microbial community experimentally, the beetles suffered a significant, though sub-lethal reduction in fitness. The design of our experiment means we can attribute these effects directly to the microbial community on the carcass, rather than to the community of microbes carried by beetles within their gut or in the surrounding soil.

At first sight, our results seem surprising because previous work has shown that the microbes associated with carrion can reduce burying beetle reproductive success and larval growth (Rozen et al 2008). Furthermore, burying beetles actively avoid carcasses that carry very high concentrations of microbes (Rozen et al 2008). Breeding on germ-free mice therefore seems more likely to promote beetle reproductive success than to reduce it. However, burying beetles produce antimicrobial exudates (Cotter & Kilner 2010, Palmer et al 2016), which eliminate some members of the microbial community on the carcass (e.g. Duarte et al 2017). Furthermore, burying beetle larvae grow and survive better on carcasses that have been treated with these exudates than on those which have not (Rozen et al 2008). Our results show that the microbes that remain on the carcass, and specifically those which derive from the carcass itself (Duarte et al 2017), are somehow beneficial to the beetle. In addition, the microbiota found on the carcass skin and hair, and those found in the lower gut appear to independently affect the mass attained by larvae during development (Fig 1).

Exactly how this is achieved has yet to be discovered. One possibility is that some, or all, of the microbes derived from the mouse directly assist larvae in digesting the carcass (cf Ruokolainen et al 2016). Alternatively, or as well, the microbes may induce parents to provision their offspring more extensively. We have previously shown that an experimental increase in the bacterial concentration on a carcass can induce greater levels of some components of parental care (Cotter et al 2010), but whether this applies to all aspects of care is unknown.

More generally our work shows that animals can evolve a beneficial association with a microbial community, even if that community is external to the animal, is transiently associated with it at each generation, and is not vertically transmitted directly from parent to offspring. Our data suggest that this beneficial association could arise in the first instance simply as an adaptation to the microbial ecosystem in which the animal lives (see also Douglas & Werren 2015). Recurrent exposure to this ecosystem is therefore a key initial step in the evolution of this association, and it need not involve direct vertical transmission. The next step, suggested by the burying beetle’s natural history, is that the animal evolves the ability to eliminate from the community any microbes that are detrimental to animal fitness. This could involve the animal repurposing existing immune defences to produce antimicrobial secretions, for example (e.g. Palmer et al 2016). Management of the microbial ecosystem in this way can then pave the way for the animal to derive fitness benefits from the microbes that remain. Internalisation of the microbial community, and direct methods for its vertical transmission from parents to offspring, may follow later. But, our results suggest, it is the animal adapting to its microbial ecosystem that sets this cascade of evolutionary events in motion.

## SUPPLEMENTARY INFORMATION

*Livefoods conventional mice versus HsdOla:MF1 conventional mice*

## METHODS

It is theoretically possible that our laboratory population of burying beetles was locally adapted to conventional albino mice, independent of any affects of the associated microbial community. To test this possibility, we also bred beetles on HsdOla:MF1 mice (i.e. the same stain of mice as the germ-free mice, obtained frozen and dead from the Gurdon Institute and Biomedical Facility at Cambridge University, and compared their performance with beetles breeding on albino white mice. Once again, we cut mice into head (LiveFoods Direct mouse 18.50 ± 0.31 g, HsdOla:MF1 strain 18.57 ± 0.34 g) and tail portions (LiveFoods Direct mouse 17.43 ± 0.38 g, HsdOla:MF1 strain 17.40 ± 0.37 g) as described above.

## RESULTS

We found no difference in the overall breeding success of beetles on LiveFoods Direct albino conventional mice versus HsdOla:MF1 conventional mice: 20 out of 20 pairs on LiveFoods Direct mice bred successfully compared with 19 out of 20 pairs on HsdOla:MF1 mice (GLM with binomial distribution, logit link function; *X^2^* = 0.00, *p* = 1.00). We found that beetles that bred on the HsdOla:MF1 mice produced a larger clutch of offspring than those on the LiveFoods Direct mice (LM; F_1, 34_ = 8.97, *p* = 0.005), and that this was irrespective of the portion of the carcass on which they developed (LM; F_1, 34_ = 0.002, *p* = 0.96). Consequently, we found that the mass of the broods on the HsdOla:MF1 mice were larger than on the LiveFoods Direct mice (LM; F_1, 34_ = 9.43, *p* = 0.004) and that this was irrespective of the portion of the carcass on which they developed (LM; F_1, 34_ = 0.004, *p* = 0.95). However, after controlling for larval density on the carcass (Table S1), we found no significant difference in the average mass of larvae produced between treatments (Fig. S1a, b, Table S2) nor carcass portion (Fig. S1b, Table S2). Therefore, we find no evidence that the burying beetles are locally adapted to the strain of mouse *per se* that we breed them on in the laboratory.

## Supplementary Figures

**Figure S1.**
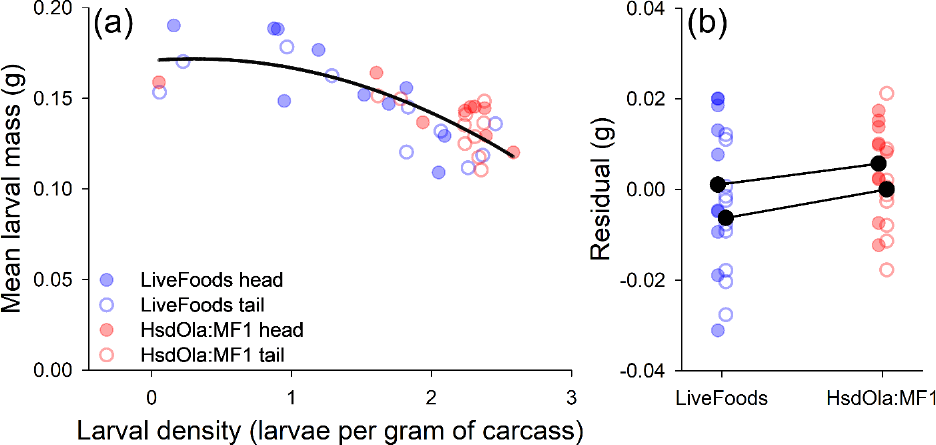
The mean larval mass of members of the brood at dispersal. A) A quadratic regression (*R^2^* = 0.60) describes the relationship between mean larval mass and larval density. B) The relative size of larvae at dispersal do not significantly differ according to carcass type, carcass part, or their interaction.

## Supplementary Tables

**Table S1.** Best-fit regression model of mean larval mass on larval density for all broods produced on LiveFoods Direct and HsdOla:MF1 mice.

| Source | *Estimate* | *Std error* | *t* | *P* | *R*<sup>2</sup> |
| --- | --- | --- | --- | --- | --- |
| Intercept | 0.2027 | 0.0105 | **19.37** | **<0.001** | 0.60 |
| Larval density | -0.0300 | 0.0047 | **-6.32** | **<0.001** |  |
| Larval density<sup>2</sup> | -0.0104 | 0.0045 | **-2.29** | **0.028** |  |

**Table S2.** A linear model testing for the effect of carcass treatment, carcass part, and their interaction on the relative size of larvae at dispersal that developed on LiveFoods Direct or HsdOla:MF1 mice.

| Source | *Estimate* | *Std error* | *t* | *P* |
| --- | --- | --- | --- | --- |
| Intercept | 0.0001 | 0.0022 | 0.07 | 0.947 |
| Treatment (LiveFoods) | -0.0027 | 0.0022 | -1.27 | 0.212 |
| Part (Head) | 0.0033 | 0.0022 | 1.51 | 0.139 |
| Treatment (LiveFoods) × Part (head) | 0.0004 | 0.0022 | 0.19 | 0.851 |

